# Preserved perception-action dissociation but altered visuomotor behaviours in healthy aging

**DOI:** 10.1101/2025.10.27.684787

**Authors:** Felicia Tassone, Zoha Ahmad, Tzvi Ganel, Erez Freud

## Abstract

The two visual pathways hypothesis posits distinct brain systems for vision-for-perception and vision-for-action. While this dissociation is well-established in younger adults, its integrity in healthy aging remains unclear. To address this, younger (*n* = 25, range: 18–25 years) and older adults (*n* = 25, range: 65–95 years) completed estimation and grasping tasks in two experiments. In Experiment 1, two rectangular objects with varying lengths (40 mm and 42 mm) were placed on the “far” and “close” surfaces of a Ponzo illusion. Despite age-related changes in grasping kinematics, the perception–action dissociation persisted. The Ponzo illusion influenced estimation such that objects placed on the “far” surface were perceived as longer. In contrast, grasping was not affected by the illusion, and even showed effect in the opposite direction, with larger apertures for objects placed on the “close” surface of the illusion. Experiment 2 tested whether this reversed effect was mediated by the surface size on which the object was placed, rather than perceived distance. To this end, we removed the illusory distance cues and varied only the background surface size (“big” versus “small”). While perceptual estimations were unaffected, surface size modulated grasping in both age groups, with a stronger effect in older adults. These findings indicate that the perception–action dissociation is preserved in aging, but older adults rely more on contextual cues during action, potentially reflecting compensatory mechanisms to maintain visuomotor performance.

## 1. Introduction

The dissociation between visual perception and action represents a fundamental organizing principle in cognitive neuroscience. According to this framework, visual processing diverges into two anatomically and functionally distinct streams: the ventral pathway supporting object identification and recognition (“vision-for-perception”) and the dorsal pathway facilitating visually guided actions (“vision-for-action”) (Goodale & Milner, 1992). Crucially, this dissociation is not a fixed, monolithic property of the visual system, but rather dynamically emerges through development (e.g., Freud et al., 2019; Hadad et al., 2012; van der Kamp & Savelsbergh, 2000), varies across individuals, and may be altered by aging-related neural changes. Understanding how this fundamental dissociation manifests across the lifespan could provide critical insights into the flexibility and constraints of visuomotor specialization.

While initial studies on the dissociation focused on neuropsychological cases (e.g., Goodale et al., 1991), visual illusions have emerged as powerful experimental tools for revealing the functional independence of perception and action systems in neurotypical adults. Illusions such as the Ponzo, Ebbinghaüs/Titchener, and Müller-Lyer typically induce robust perceptual biases: participants consistently misjudge object sizes or lengths based on contextual surroundings. However, when participants reach to grasp these same objects, their grip aperture often scales appropriately to the actual size, remaining largely unaffected by the illusory context (Aglioti et al., 1995; Ganel et al., 2008b; Ozana & Ganel, 2020). This behavioural divergence demonstrates that perception and action rely on qualitatively different visual computations: perceptual estimates are systematically biased by contextual information, while grasping apertures scale to actual object properties, despite the illusory context. This interpretation is strengthened by other paradigms that show that perceptual judgments typically follow Weber’s law, while grasping movements often violate this principle, suggesting action relies on absolute rather than relative metrics (Ganel et al., 2008a).

Nevertheless, the comparison between perceptual and visuomotor performance has faced important criticisms. Some researchers argue that differences between perception and action tasks (such as feedback availability, temporal demands, and required spatial precision) rather than distinct neural representations or distinctive processing styles may account for differential illusion effects (Franz, 2001; Franz & Gegenfurtner, 2008; Smeets & Brenner, 2006). Importantly, although such concerns have been primarily raised in the case of specific illusions, such as the Ebbinghaus, there are other illusions–in particular, the Ponzo illusion–that show consistent dissociations between action and perception, regardless of feedback or other experimental concerns. Moreover, evidence from specific populations, for example, children with amblyopia (Ahmad et al., 2023), shows that these groups remain sensitive to visual illusions in both perceptual and grasping tasks. Such findings reinforce that illusion paradigms can indeed provide a powerful window into the perception–action dissociation, even in contexts where alternative explanations have been raised.

In typically developing children, the perception-action dissociation is present from infancy, but continues to mature throughout childhood. Research indicates that this dissociation may already be present quite early in infancy: 6-to 7-month-old infants can perceive depth illusions and are influenced by allocentric cues in their perception, but their reaching movements are primarily guided by physical object distance, not perceptual illusions (van Wermeskerken et al., 2012). This dissociation strengthens between ages 5-8, when children’s grasping movements consistently resist visual illusions despite their perceptual judgments remaining strongly biased (Freud et al., 2021). This developmental trajectory suggests that the visual systems for perception and action develop as partly separate pathways from early in life, supporting the two-stream model of visual processing (Goodale & Milner, 1992; Van Der Kamp, J. & Savelsbergh, 2000).

However, the developmental trajectory itself is quite complex. In early childhood (18 to 30 months) children perform more scaling errors, such as failing to use real-world information about object size and attempting to perform impossible actions on miniature objects (e.g., attempting to sit in a tiny car, as seen in DeLoache et al., 2004). These errors in scaling may reflect a greater integration between perceptual and motor processes early in life. Additionally, children consistently follow Weber’s law in both in perceptual tasks and in grasping for complex, but not for simple objects, whereas adults show consistent violations of Weber’s law in grasping, regardless of object complexity (Freud et al., 2019; Hadad et al., 2012). These findings suggest that perception and action are more tightly coupled in early childhood, with functional segregation strengthening throughout development. Moreover, as discussed above, atypical developmental conditions reveal the dissociation’s vulnerability to altered neural organization. Individuals with autism spectrum disorder, Williams syndrome, amblyopia, and resection of both cortical pathways often show reduced or absent perception-action dissociation, indicating atypical dorsal stream processing or altered integration between visual pathways (Ahmad et al., 2022; Ahmad et al., 2023; Ahmad et al., 2025; Dilks et al., 2008). These findings collectively demonstrate that the perception-action dissociation is experience-dependent and malleable, raising important questions about its stability during aging-related neural reorganization. Additional findings suggest that the dissociation between perception and action for size illusions can be compromised even in typically developed adults. In particular, grasping movements in left-handed–but not in right-handed–participants are affected by the Ponzo illusion when grasping with the right, non-dominant hand (Ganel & Goodale, 2024; but also see Gonzalez et al., 2006).

Among visual illusions used to study perception-action dissociation, the Ponzo illusion offers unique methodological advantages that make it particularly suitable for examining age-related changes. The illusion employs converging lines to simulate depth perspective, causing identical objects placed at different positions to appear dramatically different in size perceptually: objects at the “far” position appear longer than those at the “close” position, and the perceptual effect is consistent across age-groups (e.g., Mazuz et al., 2024). Previous research in young adults has consistently demonstrated that while perceptual estimates show strong susceptibility to the Ponzo illusion, grasping movements remain largely veridical, scaling to actual rather than perceived size (Ganel et al., 2008b; Ozana & Ganel, 2020; Whitwell et al., 2016). However, the dissociation is not absolute: when participants are asked to stop their grasp at maximum grip aperture rather than completing the movement (before object contact), their grip becomes biased by the Ponzo illusion (Navon & Ganel, 2020). This suggests that conscious monitoring of movement can introduce perceptual intrusions into otherwise veridical grasping. The reliability and magnitude of this dissociation, combined with its intuitive depth-based mechanism, positions the Ponzo illusion as a powerful probe for investigating how aging might alter the functional separation between perception and action systems.

Aging brings profound changes to visual, motor, and cognitive systems that could fundamentally alter the perception-action dissociation (Grady, 2012). This decline is marked by a pronounced deterioration of various visual behaviours, including reduced visual acuity, depth perception, and contrast sensitivity (Song et al., 2023; Swenor et al., 2018). Motor behaviour also becomes less efficient: older adults typically show slower movement execution, larger grip apertures, and reduced grasping precision (Campoi et al., 2023; Maki & McIlroy, 2006; Vasylenko et al., 2018; Voelcker-Rehage & Alberts, 2005). These changes may reflect not only physical constraints, but also neural shifts in brain structure and function, including reduced grey matter volume and altered functional connectivity (Carp et al., 2011; Park et al., 2004; Raz et al., 2005). Nevertheless, the effect of aging on the perception-action dissociation is largely unknown and is the focus of the current investigation.

Two major theoretical frameworks offer competing predictions. The dedifferentiation hypothesis proposes that aging reduces neural specialization, leading to increased activation overlap and decreased functional segregation between brain systems (Koen & Rugg, 2019). Neuroimaging evidence supports this model: older adults show reduced distinctiveness of neural representations in both visual ventral cortex and motor control networks, suggesting that previously specialized systems become increasingly integrated (Carp et al., 2011; Park et al., 2004). Under this framework, the boundaries between dorsal and ventral visual pathways would blur with age, predicting that older adults’ actions would become increasingly susceptible to perceptual illusions; effectively weakening or eliminating the perception-action dissociation. Alternatively, the compensation-related utilization of neural circuits hypothesis (CRUNCH) suggests that older adults recruit additional or alternative brain regions to maintain performance despite age-related neural decline (Reuter-Lorenz & Cappell, 2008). This compensatory recruitment might preserve the functional dissociation between perception and action while altering the specific strategies used to achieve accurate performance. Supporting this view, older adults show increased reliance on external visual cues and feedback during reaching movements, potentially indicating adaptive compensatory mechanisms (Wermelinger et al., 2018). In the current study, we examine which of these models can better account the effect of aging on the perception-action dissociation. While the dedifferentiation model predicts degraded dissociation with increased illusion susceptibility in both tasks, the CRUNCH model predicts preserved dissociation maintained through compensatory strategies (Figure 1).

**Figure 1.**
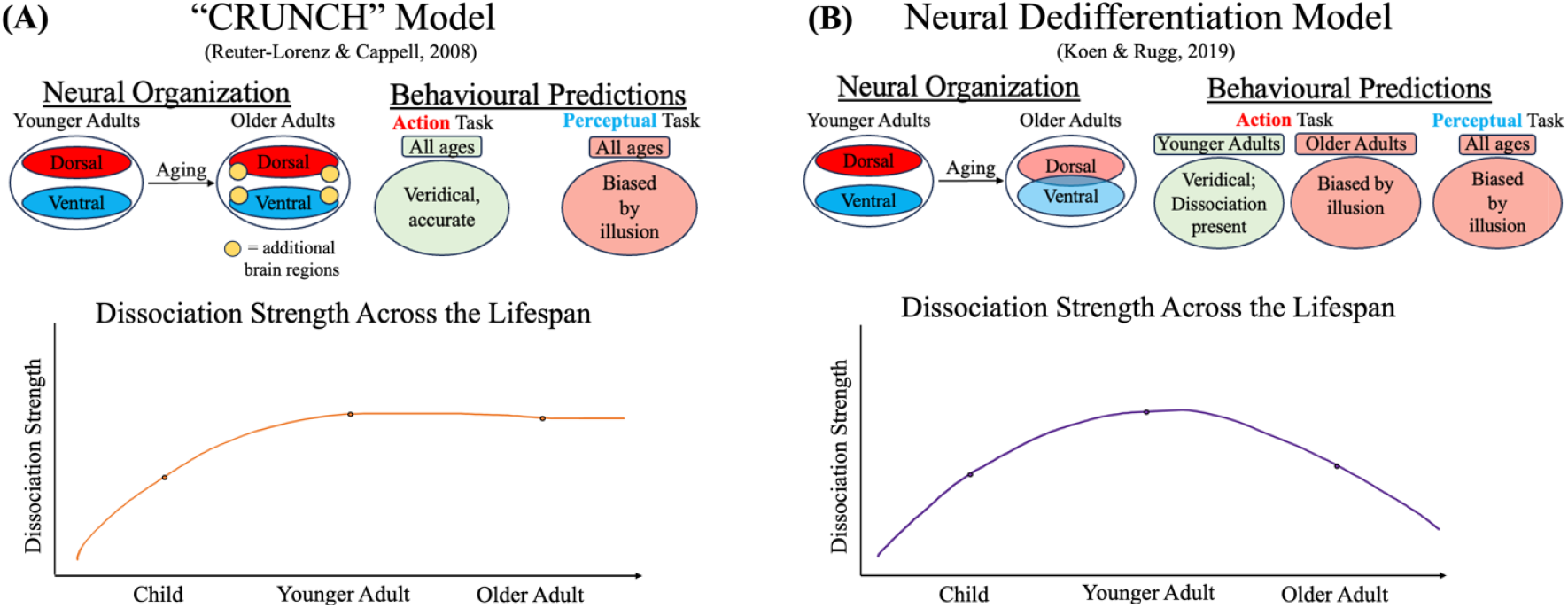
Schematic predictions of the*”CRUNCH” Model (Reuter-Lorenz & Cappell, 2008)* **(A)** and the *Neural Dedifferentiation Model (Koen & Rugg, 2019)* **(B)** for the relations between perception and action across the lifespan.

## 2. Experiment 1

### 2.1. Methods

#### 2.1.1. Participants

Twenty-five younger adults (14 female; *M* = 19.88 years, *SD* = 1.81, range = 18–25) and twenty-five older adults (23 female; *M* = 76.04 years, *SD* = 5.30, range = 69–95) participated in Experiment 1. Sample size was determined based on previous studies in which the perception-action dissociation was compared as a between-groups variable (Ahmad et al., 2023; Ahmad et al., 2025; Freud et al., 2021). All participants self-identified as right-handed, confirmed using a modified version of the Edinburgh (Oldfield, 1971) and Waterloo (Brown et al., 2006) handedness questionnaires (see Stone et al., 2013 for details). Visual acuity was verified using a Snellen chart. Younger adults were recruited from York University’s undergraduate research pool, while older adults were recruited from the Elspeth Hayworth Centre for Women. Older adults were healthy and independent enough to arrive to the centre and participate in weekly activities and classes.

The study was approved by the Research Ethics Board of York University. All participants provided informed, signed consent prior to testing, after receiving an explanation of the study’s procedures and potential risks. Participants received either course credit (younger adults) or monetary compensation (older adults). The procedures were based on methodologies used in previous work on grasping and perceptual estimation under visual illusions (e.g., Ahmad et al., 2022; Freud et al., 2021).

#### 2.1.2. Apparatus and Stimuli

Participants sat in front of a table on which a printed Ponzo illusion board was placed (Figure 2A). The board consisted of a flat surface containing converging diagonal lines that created illusory depth. Objects were positioned at either the “close” or “far” surface of the board. Two black rectangular 3D plastic blocks were used as target objects, matched in height (10 mm), width (10 mm), and color (black), but differing slightly in length: one measured 40 mm (“shorter”) and the other 42 mm (“longer”).

**Figure 2.**
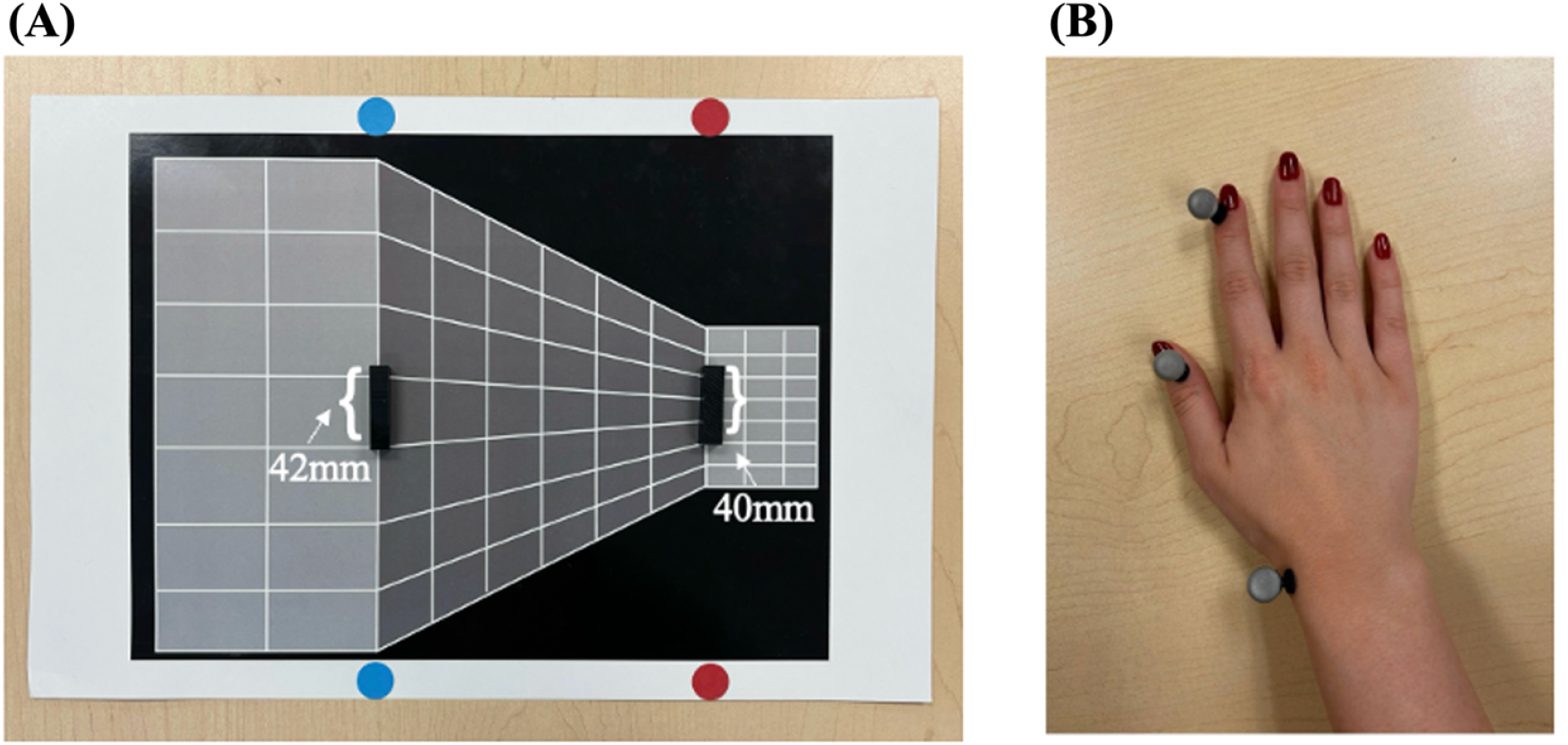
**(A)**. *Experimental setup*. The “close” surface is marked with blue stickers and the “far” surface is marked with red stickers. Note that both physical objects were placed on a flat 2d background of the illusion. Due to depth cues, objects placed on the far surface are perceived to be longer than they are in actuality. **(B)**. *Marker placement*. Reflective markers placed on the index finger, thumb, and wrist.

Kinematic data were recorded using the OptiTrack motion-capture system at a sampling rate of 100 Hz. Five Prime 13w cameras captured participants’ grasping and estimation movements at x (left-right), y (up-down), and z (posterior-anterior) coordinates. Three reflective spherical markers were attached to the participant’s index finger, thumb, and wrist (Figure 2B).

#### 2.1.3. Procedure

Participants completed two tasks: a grasping task and a perceptual estimation task, with the order counterbalanced across participants.

Each trial began with participants at a “home base” position, with their thumb and index finger together. Two identical objects were manually placed simultaneously: one on the close surface, and one on the far surface. In each task, each object (shorter and longer) appeared on either the close or far surfaces, for a total of 30 randomized presentations per condition (object size × location), resulting in 60 trials per task. The experimenter manually initiated each trial. After a 1-second delay, an auditory cue (“BLUE” or “RED”) was generated by the computer. “BLUE” indicated the object on the close surface, while “RED” indicated the object on the far surface. Participants initiated the grasping or estimation movement and had no time constraints to complete their response. The Ponzo board was rotated every 10 trials to ensure that the “close” and “far” surfaces are not associated with left or right movements.

In the grasping task, participants reached for the cued object, grasped it along its length using their thumb and index finger, lifted it, placed it back at its original position, and returned to the home base with thumb and index finger together. In the estimation task, participants moved their hand to their right side, opened their thumb and index finger to indicate the perceived length of the cued object, held this aperture for approximately 1 second, and then returned to the home base with their thumb and index finger together.

#### 2.1.4. Data Analysis

Data were processed and analyzed using custom, in-house Python scripts and JASP (JASP Team, 2024). For grasping trials, 3D trajectories of the object, index finger, thumb, and wrist were recorded. Grip aperture was calculated as the Euclidean distance between the thumb and index finger markers. Movement onset was defined as the first frame at which either the index finger or wrist velocity exceeded 20 mm/s for 20 consecutive frames. Movement offset was determined using two algorithms: (1) a velocity-based condition in which the object velocity exceeded 50 mm/s while aperture change remained under 0.5 mm for 10 consecutive frames, and (2) a position-based condition where grasp completion was identified by marker position thresholds in the Y and Z axes combined with low aperture variability. The earliest valid time point across the two criteria was selected. For estimation trials, movement onset was defined using the same velocity threshold as in grasping, while movement offset was marked by the final frame of the recording (“endpoint”). All trials were visually inspected and manual adjustments were made when the algorithm failed to detect clear movement boundaries.

For each grasping trial, grip aperture values were extracted at every frame from movement onset to movement offset. The maximum grip aperture (MGA) was identified as the peak aperture between movement onset and offset. For estimation trials, the aperture value at the final frame was used to estimate perceived object length, as participants did not physically interact with the object.

To correct for individual variation in finger width (which might induce artificial aperture differences between age groups), MGA values were normalized using a procedure adapted from Ganel et al. (2012). For each participant, the grip aperture at the final frame of each trial was averaged across all trials. Then, a constant value of 40 mm (corresponding to the length of the shorter object) was subtracted from this average to estimate each participant’s finger width. Note that subtracting the fixed value of the shorter object rather than the actual object size ensured that effects of size were preserved. This estimated finger width was then subtracted from all MGA values and from the grip aperture values at each time point (0–100%) for every trial. This correction could not be applied to estimation trials since the endpoint of the trial does not reflect the true size of the objects, so analyses for that task were conducted on raw values.

Effect of the illusion was calculated by subtracting the average response for “close” trials from “far” trials, separately for each object size and task. Positive values reflected the expected influence of the Ponzo illusion, wherein the distance between fingers is greater for objects placed on the “far” surface. Size sensitivity was calculated as the difference in response between big and small objects, collapsing across object’s location. Group comparisons and task effects were assessed using ANOVAs, planned simple comparisons, and t-tests.

## 3. Results

### 3.1. Perception-Action Dissociation

Figure 3A presents the effect of the illusion on grasping and on perceptual estimations for the two age groups. As can be seen in the figure, perceptual estimations, but not grasping, were susceptible to the illusion for both age groups. To examine whether the Ponzo illusion differentially influenced perception and action, a mixed-design ANOVA with task, age-group, and perceived distance was conducted on endpoint (perceptual task) and adjusted MGA values (action task). This analysis revealed a robust interaction between task and perceived distance [*F*_(1, 48)_ = 44.99, *p* <.001, *η*^*2*^*p* =.484], with no interaction between age-group and the other two factors [*F*<1], suggesting that the dissociation was preserved in both age groups (Figure 3A). This was confirmed by follow-up ANOVAs conducted separately for each age group that showed significant interactions between task and perceived distance [Younger: *F*_(1, 24)_ = 24.52, *p* <.001, *η*^*2*^*p* =.505; Older: *F*_(1, 24)_ = 22.18, *p* <.001, *η*^*2*^*p* =.480].

**Figure 3.**
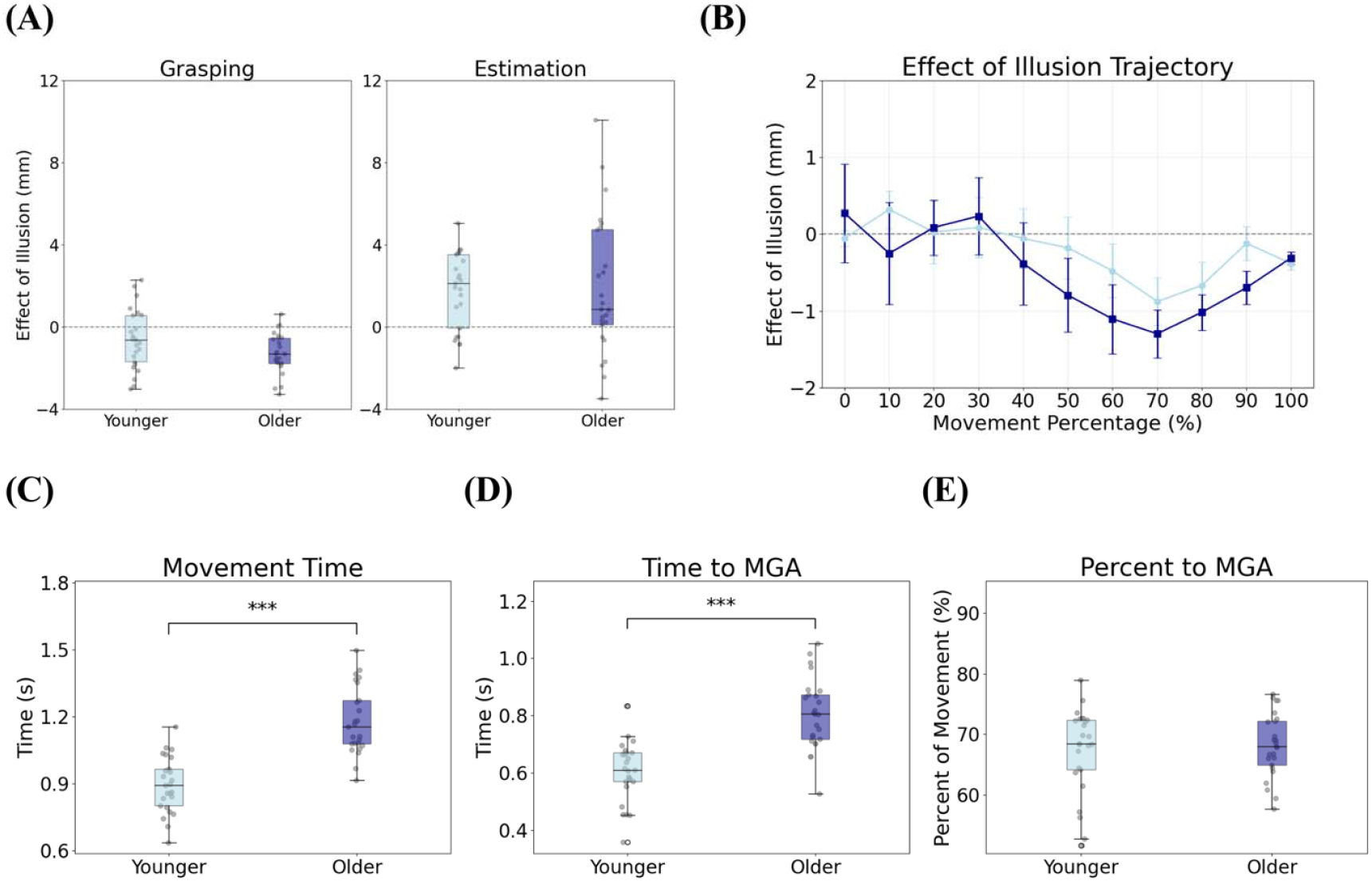
Results for Experiment 1. **(A)**. Illusory effects for grasping (left) and estimation (right) across groups, derived from finger aperture in millimetres. Boxplots show group-level distributions of the illusory effect (far – close), with individual data points overlaid. **(B)**. Effect of the illusion (far – close) plotted across normalized movement time, calculated at 10% intervals from movement onset to offset. **(C)**. Movement time (in seconds) by group. (**D)**. Time to MGA (in seconds) by group. **(E)**. Time to MGA expressed in terms of percentage of the movement. Error bars in all figures represent standard error of the mean (SE).

As expected, perceptual estimates were significantly modulated by the illusion, with objects appearing larger when placed on the far surface compared to the close surface [*F*_(1, 48)_ = 23.54, *p* <.001, *η*^*2*^*p* =.329]. Planned comparisons revealed that this was true for both younger [*t*(48) = 3.30, *p* =.002] and older adults [*t*(48) = 3.57, *p* <.001]. The effect of the illusion was similar across the two age groups [*F*_(1, 48)_ = 0.037, *p* =.849].

Unexpectedly, grasping responses showed a small but reliable effect in the opposite direction: grip apertures were larger for objects on the close surface compared to the far surface [*F*_(1, 48)_ = 29.52, *p* <.001, *η*^*2*^*p* =.381]. Planned comparisons revealed this reversed effect was significant in both younger adults [*t*(48) = –2.49, *p* =.016] and older adults [*t*(48) = –5.19, *p* <.001], with means of –0.62 mm (*SD* = 1.44) and–1.28 mm (*SD* = 0.98), respectively. The interaction between group and perceived distance approached significance [*F*(_1, 48)_ = 3.64, *p* =.062, *η*^*2*^*p* =.071], suggesting a trend toward a larger reversed effect in older adults. This unexpected finding motivated Experiment 2, where we examined whether this reversal was specific to the Ponzo illusion context.

Despite the observed perception-action dissociation observed for both age groups, it remains possible that older adults’ grasping behaviours might be modulated by the illusion at other stages of the movement trajectory (i.e., before or after the MGA). To test this idea, we computed the illusion effect at each 10% interval (far – close) (Figure 3B). Both age groups showed a negative illusion effect during mid-to-late stages of movement, consistent with the reversed MGA pattern. No significant group differences were found at any timepoint (*ts* < 1.88; *ps* >.723), suggesting that both groups were similarly affected by the illusion across the movement trajectory.

### 3.2. Age-Related Kinematic Variables

Although the two groups exhibit a similar pattern of perception-action dissociation, older adults exhibited significant kinematic differences, suggesting altered visuomotor execution. We compared the temporal profile of the grasping movements using an independent t-test that revealed that older adults had longer movement time [*t(48)* = 7.32, *p* = <.001, *d* = 0.407] and took longer to reach the MGA [*t(48)* = 6.26, *p* = <.001, *d* = 0.378] (Figures 3C and 3D).

Consistent with previous literature (Ganel et al., 2008b; Ozana & Ganel, 2020), these results suggest that although grasping responses were largely resistant to the illusory context in terms of aperture scaling, the temporal dynamics of movement execution varied with age. Yet, despite these visuomotor differences when looking at absolute timing, the proportional structure of the movement (measured by percent time to MGA) did not differ between groups (*t* = 0.44, *p* =.662) (Figure 3E). This suggests that although older adults moved more slowly and reached peak grip aperture later in time, the relative timing within the reach was preserved.

### 3.3. Interim Discussion

Experiment 1 confirmed perception-action dissociation in both age groups, consistent with preserved functional segregation. As predicted by the two visual pathways hypothesis (Goodale & Milner, 1992), participants exhibited a strong susceptibility to the Ponzo illusion during estimation trials, wherein objects placed on the “far” surface were consistently judged to be larger, while this effect was not observed during the grasping task. However, the present data also revealed a surprising reversal in the direction of the effect during grasping: both younger and older participants opened their fingers wider for objects located on the “*close”* surface compared to the *“far”* surface. This reversed effect was greater for older adults.

These results suggest that participants’ grasping behaviour was influenced by some component of the visual display. One plausible explanation for this reversal effect lies in the relative size of the surfaces in the Ponzo illusion board. In particular, the “close” surface in the Ponzo display is physically larger than the “far” surface (Figure 2A). Thus, it is possible that grasping movements were influenced not by the illusory depth per se, but by the visual properties of the surface immediately surrounding the object. For example, the larger size of the close surface may have altered the perceived affordances of the grasping environment, thereby prompting participants to scale their grip differently than we anticipated (Fagg & Arbib, 1998).

Moreover, this effect may be especially pronounced in older adults due to age-related changes in visuomotor integration and strategy selection. Previous research has shown that older adults may exhibit more compensatory, conservative, and variable grasping strategies, often characterized by larger grip apertures and slower movements (Campoi et al., 2023; Cicerale et al., 2014). The data from our study further support this pattern, with older adults demonstrating slower movement times and longer times to peak grip aperture relative to younger adults. These motor patterns may reflect a compensatory strategy aimed at minimizing risk and ensuring task success. In this context, the visual layout of the close surface may have exerted an exaggerated influence on their motor planning, leading to the observed reversal effect.

To further investigate this possibility, we designed a follow-up experiment that eliminated the illusion-inducing perspective lines of the Ponzo board. Experiment 2 utilized flat, 2D printed surfaces that varied in their physical dimensions but did not generate an illusion of depth (Figure 4). These surfaces retained the essential manipulation of surface size (i.e., a large “close” surface and a small “far” surface) while removing the classic illusory depth cues present in the Ponzo illusion (see Ahsan et al., 2025 for a similar manipulation). The goal was to determine whether the reversed effect during grasping observed in Experiment 1 could be attributed to surface size alone.

**Figure 4.**
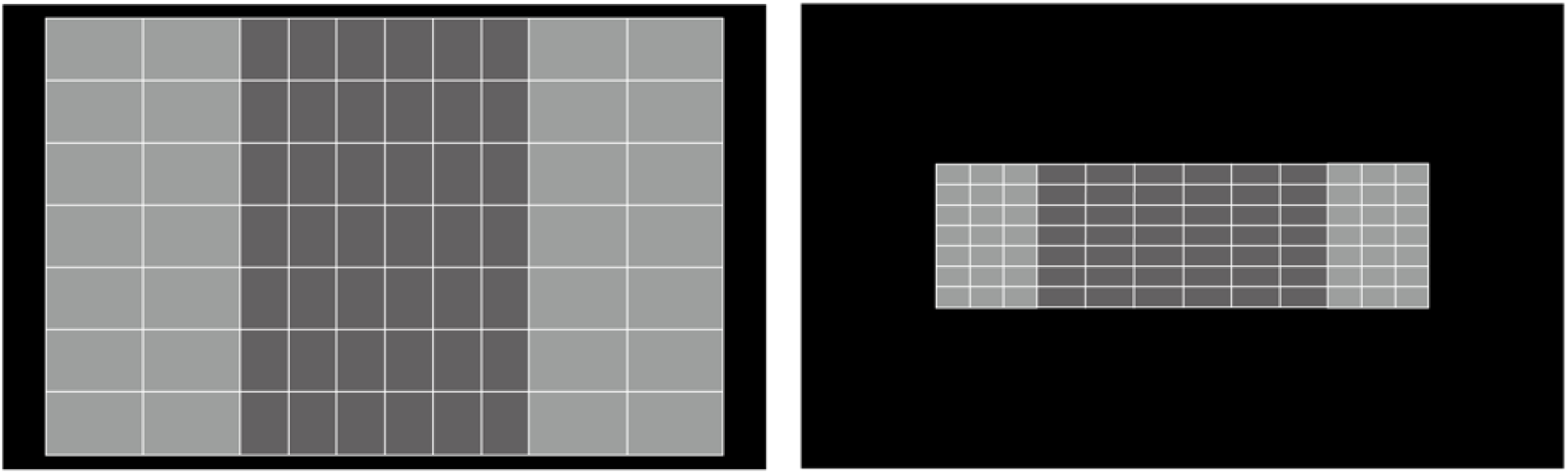
*Experimental set-up*. On the left is the “close”, *bigger* surface background, and on the right is the “far”, *smaller* surface background.

If grasping behaviour in the Ponzo illusion condition was indeed driven by visual context (e.g., background surface size) rather than by the illusion itself, then we would expect to observe a similar pattern in Experiment 2; that is, larger grip apertures for objects on the bigger (“close”) surface, even in the absence of an illusion. Conversely, if the reversed effect was attributed to other, anomalous illusion-based phenomenon, it should disappear when the illusory cues are removed.

## 4. Experiment 2

### 4.1. Methods

#### 4.1.1. Participants

Twenty-five younger adults (15 female; *M* = 20.60 years, *SD* = 2.20, range = 18–25) and twenty-five older adults (21 female; *M* = 75.80 years, *SD* = 4.76, range = 68–87) participated in Experiment 2. Participant eligibility, screening criteria, and recruitment procedures were identical to Experiment 1. All participants provided informed consent and received either course credit (younger) or monetary compensation (older). Some younger (*n* = 4) and older (*n* = 14) adults participated in both experiments, with a 10-month gap between sessions.

#### 4.1.2. Apparatus and Procedure

The apparatus and procedures mirrored those of Experiment 1, except that the illusion-inducing diagonal lines were removed. Participants performed the same grasping and estimation tasks using the same target objects (40 mm and 42 mm blocks). However, objects were presented on flat surfaces that mimicked the spatial layout of the Ponzo illusion, without inducing depth cues (Figure 4). The “close/bigger” surface was physically larger than the “far/smaller” surface, allowing us to isolate the role of surface size in modulating visuomotor behaviour.

#### 4.1.3. Design and Procedure

Each participant completed 60 grasping trials and 60 estimation trials. The task structure, randomization, and trial sequence were identical to those in Experiment 1. The order of tasks was counterbalanced across participants. In grasping trials, participants picked up the object with their index finger and thumb. In estimation trials, they extended their thumb and index finger to indicate their perceived length of the object without touching it.

#### 4.1.4. Data Analysis

Data recording, processing, movement onset/offset definitions, and normalization procedures were identical to those in Experiment 1. The only difference was the independent variable of interest: instead of computing the effect of the illusion, we computed the effect of surface size, measured as the difference between “small” and “big” trials. Positive values indicated larger responses for objects on the small surface. Statistical analyses (ANOVAs and t-tests) examined effects of surface size, object size, and group.

## 5. Results

### 5.1. Perception-Action Dissociation

The effect of surface size on apertures during grasping and perceptual estimations is presented in Figure 5A. As can be seen in the figure, grasping apertures in both age groups showed an effect of context. Perceptual estimations, on the other hand, did not show this effect. To examine whether surface size influenced perception and action differently, a mixed-design ANOVA with task, surface size, and group was conducted on adjusted MGA values and estimation values. This analysis revealed a significant interaction between task and surface size [*F*_(1, 48)_ = 20.53, *p* <.001, *η*^*2*^*p* =.300], indicating a dissociation: grasping was modulated by surface size [*t*(95.73) = 4.38, *p* <.001], whereas estimation was not significantly modulated, but nears significance in the reversed direction [*t*(95.73) = –1.86, *p* =.066] (Figure 5A). There was no significant interaction between group, task, and surface size [*F*_(1, 48)_ = 2.55, *p* =.117], suggesting that the dissociation pattern held across age groups.

**Figure 5.**
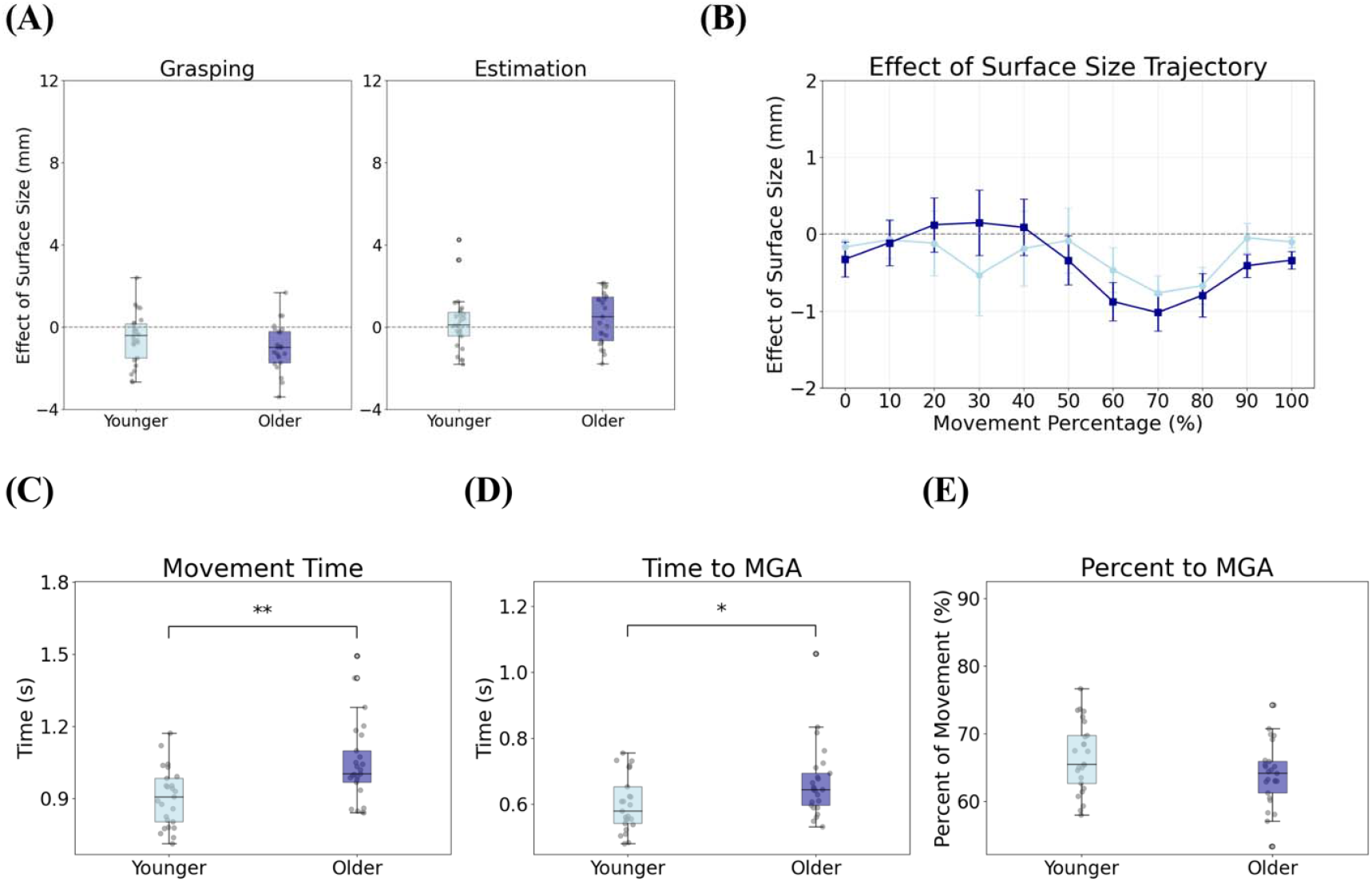
Results for Experiment 2. **(A)**. Surface size effects for grasping (left) and estimation (right), derived from finger aperture in millimetres, plotted by group. Boxplots represent the effect of surface size (small – big), with individual data points overlaid. **(B)**. Effect of surface size (small – big) plotted across movement time, calculated at 10% intervals from movement onset to offset. **(C)**. Movement time (in seconds) by group. **(D)**. Time to MGA (in seconds) by group. **(E)**. Time to MGA expressed in terms of percentage of the movement. Error bars in all figures represent standard error of the mean (SE).

Estimation responses were not significantly affected by surface size [*F*_(1, 48)_ = 3.30, *p* =.076]. At the group level, younger adults showed no effect [*t*(48) = 0.85, *p* =.400], while older adults showed a non-significant trend toward an effect [*t(*48) = 1.72, *p* =.092].

Importantly, the focus of Experiment 2 was the effect of surface size of grasping behaviors. Indeed, grasping responses were significantly affected by this factor: participants opened their fingers wider when grasping objects placed on the big surface in comparison to the small surface [*F*_(1,48)_ = 20.06, *p* <.001, *η*^*2*^*p* = 0.295]. This surface size effect was evident in younger adults [*t*(48) = –2.03, *p* =.048] and in older adults [*t*(48) = 4.30, *p* = <.001]. The interaction between group and perceived distance was not significant [*F*_(1,48)_ = 2.58, *p* =.115].

To further assess how surface size influenced grasping dynamics, we computed the surface size effect (big – small) at each 10% interval from movement onset to offset (Figure 5B). Both age groups showed negative values at mid-to-late movement stages, indicating consistently larger grip apertures for objects on the close surface. No significant group differences were found at any timepoint (all *ps* >.05), reinforcing that both age groups exhibited similar context-related modulation across the reach trajectory.

### 5.2. Age-Related Kinematic Variables

Similar to the results of Experiment 1, older adults exhibited several motor differences from younger adults. We compared the temporal profile of the grasping movements using an independent t-test that revealed that older adults had longer movement time (*t* = 3.42, *p* =.001, *d* = 0.314) and took longer to reach the MGA (*t* = 2.37, *p* =.022, *d* = 0.298) (Figures 5C and 5D).

Percent time to MGA, which reflects the proportional temporal structure of the reach, did not reach statistical significance but showed a trend toward group differences (*t* = –1.88, *p* =.066) (Figure 5E). Unlike as in Experiment 1 where this measure showed no group differences, Experiment 2 revealed a marginal trend, suggesting that older adults may reach their MGA slightly later along the movement trajectory. However, this trend indicates that although older adults may have moved more slowly in absolute terms, the relative timing of their grasp formation was largely preserved.

### 5.3. Age Differences Across Both Experiments

Across both experiments, we observed that older adults consistently showed enhanced reversed grasping effects compared to younger adults (opening their fingers wider for objects on the “close/big” surface relative to the “far/small” surface). To test whether this age-related enhancement was consistent across experimental conditions, we conducted a two-way ANOVA (Group × Experiment) on adjusted MGA values from the grasping task.

A significant main effect of group emerged [*F*_(1, 96)_ = 6.01, *p* =.016, *η*^*2*^*p* =.059], confirming that older adults exhibited stronger reversed grasping effects than younger adults when collapsed across experiments. Importantly, there was no significant interaction between group and experiment [*F*_(1, 96)_ = 0.05, *p* =.825], indicating that this age-related enhancement was comparable whether the context involved illusory depth cues (Experiment 1) or physical surface size differences (Experiment 2).

These findings suggest that older adults are consistently more sensitive to visual context during grasping, regardless of whether that context is illusory or veridical. This pattern aligns with proposals that older adults may rely more heavily on contextual information during visuomotor control, potentially as a compensatory strategy to maintain motor accuracy in the face of age-related sensorimotor changes.

## 6. Discussion

The present study investigated the effect of healthy aging on the perception–action dissociation, as posited by the two visual pathways hypothesis (Goodale & Milner, 1992). Across two experiments, we assessed how illusory and contextual visual information influenced perception (manual estimation) and action (grasping) in younger and older adults. Across Experiments 1 and 2, we observed a consistent perception–action dissociation across age-groups, as well as distinct age-related differences in visuomotor behaviour and sensitivity to contextual information.

In Experiment 1, perception was biased, such that objects placed on the “far” surface were perceived as longer than identical objects placed on the “close” surface. The effect of the illusion on perceptual estimation was similar across age groups (for similar results, see Mazuz et al., 2024). Grasping apertures showed a reversed effect, such that participants opened their fingers wider when grasping objects on the “close” surface. This was true for both age groups, with a larger effect for the older adults. In Experiment 2, we tested whether this reversed effect could be explained by physical surface size rather than illusory depth cues. Indeed, when only surface size was manipulated, grasping responses again showed this modulated effect, with larger grip apertures for objects placed on the “big” surface rather than the “small” surface (particularly in older adults). Estimation responses were unaffected, with a non-significant trend in the opposite direction (smaller estimates for the “big” surface, *p* =.066). Together, these results provide strong behavioural evidence that perception and action are functionally dissociable in both younger and older adults.

### 6.1. Preserved Perception-Action Dissociation in Older Adults

The current study is, to our knowledge, the first to document a preserved perception–action dissociation in healthy older adults. While previous work has consistently shown that visually guided grasping in younger adults is resistant to perceptual illusions (Aglioti et al., 1995; Ganel et al., 2008b; Ozana & Ganel, 2020), no prior study had systematically tested this phenomenon in healthy aging. The only related investigation, by Skervin et al. (2021), used a stair-climbing task in which older adults’ stepping behaviour was found to be biased by perceptual distortions, suggesting a breakdown of the dissociation under locomotor–rather than manual–conditions. In contrast, the present study’s findings reveal that in hand-based actions, the dissociation between perception and action remains robust in older age, even as overall visuomotor strategies become slower and more cautious.

Despite well-documented age-related declines in cognitive, visual, sensory, and motor functions (Cicerale et al., 2014; Grady, 2012; Swenor et al., 2019), our results demonstrate that the core functional segregation between perceptual and visuomotor processes is maintained. This stands in contrast to the dedifferentiation hypothesis, which posits that aging reduces the distinctiveness of neural representations, particularly in sensory and perceptual cortical areas (Koen & Rugg, 2019; Park et al., 2004). If dedifferentiation had weakened the distinction between perceptual and visuomotor representations, older adults would have been expected to show similar biases across both the grasping and estimation tasks in Experiment 1. Instead, both age groups displayed a clear behavioural dissociation between perception and action.

This preservation of dissociation can be understood within the framework of the CRUNCH model (Reuter-Lorenz & Cappell, 2008), which proposes that older adults recruit additional neural resources to maintain performance levels despite age-related structural decline. From this perspective, the behavioural preservation of the perception–action dissociation observed here may not reflect an absence of age-related change, but rather the success of compensatory neural mechanisms described by CRUNCH. Older adults’ slower and more deliberate movements could represent behavioural manifestations of this compensation: strategic adaptations that preserve accuracy, even as efficiency declines.

Taken together, these findings suggest that while aging is associated with widespread cognitive and neural changes, the functional distinction between perception and action is preserved–likely through compensatory recruitment of additional neural resources. Behaviourally, this compensation may manifest by slower, more deliberate visuomotor strategies, that maintain accuracy even in the face of structural decline. While conclusions cannot be drawn about the neural ventral–dorsal dissociation without direct imaging data, the patterns observed here strongly suggest that the underlying computations guiding perception and action remain separated in older adulthood.

### 6.2. Sensitivity of Grasping Movements to Contextual Information

A key finding of the present study is that grasping behaviours are also sensitive to contextual information. While this might initially appear to challenge the dissociation between perception and action, the effect of contextual information differed across the two tasks. Specifically, grasping behaviours showed a reversed effect, such that grip apertures were larger for objects on the physically larger, close surface. Notably, this pattern emerged even in the absence of illusory depth cues (Experiment 2), and as such, it likely reflects a visuomotor sensitivity to physical surface layout, rather than susceptibility to perceptual distortion. Such sensitivity aligns with embodied cognition frameworks that emphasize how environmental context shapes motor planning and execution (e.g., Foglia & Wilson, 2013).

One potential explanation for this effect involves the spatial proximity of visual boundaries. In both experiments, objects placed on the “close/big” surfaces were positioned farther away from the borders of the background surface, while objects on the “far/small” surfaces were closer to them. This spatial arrangement may have influenced grip scaling through affordance-based mechanisms, wherein environmental structure modulates available action possibilities (Cisek, 2007). According to the affordance competition hypothesis (Cisek, 2007), cortical mechanisms of action selection continuously weigh multiple potential actions based on affordances present in one’s visual environment. In this view, the larger visual surface in our experiment may have afforded more space for safe movement execution, leading participants to open their grip wider.

This interpretation is supported by recent work on the Ebbinghaüs illusion by Chen et al. (2021), who found that grasping is influenced not only by illusory size information, but also by obstacle-avoidance processes. When flankers were close to the target, grip aperture was modulated as if participants were avoiding nearby objects, even when the perceptual illusion remained constant. These findings indicate that apparent “illusion effects” on action can partly reflect visuomotor adjustments to spatial layout and contextual affordances. Similarly, in the present study, the observed sensitivity of grasping to surface context may reflect adaptive visuomotor calibration to the available workspace, rather than a breakdown of perception–action dissociation.

### 6.3. Altered Kinematic Behaviours in Aging

Although both age groups demonstrated a preserved perception–action dissociation, older adults exhibited notable differences in movement dynamics and contextual sensitivity. In both experiments, older adults showed longer movement times and delayed peak grip aperture. These results are consistent with previous work showing that older adults tend to adopt more cautious, variable, and conservative grasping strategies (Caetano et al., 2016; Campoi et al., 2023; Cicerale et al., 2014).

The fact that older adults were more affected by surface size suggests they may rely more heavily on background context during action planning. One possible explanation for this increased sensitivity is provided by the CRUNCH framework (Reuter-Lorenz & Cappell, 2008). In our study, older adults may have weighted contextual visual information more heavily in order to maintain motor accuracy. Notably, this occurred without a loss of the dissociation between perception and action, further supporting the idea that aging is accompanied by strategic adaptations rather than a breakdown of core processing systems. Despite the observed slowing and greater modulation by context, percent time to MGA did not differ between groups. This suggests that although the absolute timing of movements changed with age, the temporal structure of the grasping trajectory remained intact. Thus, while older adults modified their visuomotor execution strategies, the fundamental organization of movement planning appeared to be preserved.

It is important to consider that the age-related differences we observed in visuomotor behaviour may reflect changes occurring across multiple neural systems. Aging affects both the central nervous system (CNS) and peripheral nervous system (PNS) in distinct ways. While the CNS experiences changes in brain structure, neurotransmitter function, and neural connectivity, the PNS undergoes morphological changes including loss of myelinated nerve fibers, demyelination, and reduced nerve conduction velocity (Verdú et al., 2000). Peripheral nerve aging is characterized by axonal atrophy, reduced expression of myelin proteins, and decreased regenerative capacity following injury. These changes in the PNS result in decreased sensation, slower reflexes, and reduced motor precision. As such, the PNS may be particularly vulnerable to age-related changes because peripheral nerves lack the protection of the blood-brain barrier and may be more susceptible to toxic metabolites and inflammatory mediators that accumulate with age. The slower movement times and altered grasping kinematics we observed in older adults may therefore reflect the combined influence of both central processing changes and peripheral motor system deterioration.

### 6.4. Implications for Real-World Functioning

Our findings have important implications for understanding how older adults navigate their daily environments. The increased sensitivity to contextual visual information we observed may represent not only an adaptive strategy, but also a potential vulnerability. While relying more heavily on environmental cues may help compensate for age-related sensory and motor declines, it may also increase susceptibility to misleading visual information.

For instance, research has demonstrated that complex patterns on carpets, floors, or stairs constitute an extrinsic factor contributing to fall accidents in older populations (Lu et al., 2021). Additionally, patterned surfaces can cause visual discomfort and are associated with nausea and migraines (Bonato et al., 2011; Wilkins et al., 2018). Given that visual impairments are a recognized risk factor for falls (Ivers et al., 2000), these environmental factors become particularly concerning for older adults with sensory degradation (Lockhart et al., 2005), age-related vision changes, and altered visuomotor processing.

Walking on patterned surfaces may negatively impact gait, balance, and spatial orientation (McNeil & Tapp, 2015), creating challenges for environmental recognition and navigation. Our findings suggest that while older adults’ increased reliance on contextual visual cues may serve a compensatory function, it may also make them more vulnerable to misleading or irrelevant environmental information, making it difficult to effectively organize and interpret visual signals and potentially resulting in incorrect perceptions and cognitive biases.

## 7. Conclusion

The present study examined how healthy aging modulates the perception–action dissociation using a Ponzo illusion paradigm and a matched surface-size manipulation. The findings show that while the core functional dissociation remains intact in aging, older adults may rely more on environmental context to maintain action performance. Understanding these adaptive mechanisms has important implications for designing safer, more accessible environments for aging populations.

